# Genotypic characterization of the U.S. peanut core collection

**DOI:** 10.1101/2020.04.17.047019

**Authors:** Paul I. Otyama, Roshan Kulkarni, Kelly Chamberlin, Peggy Ozias-Akins, Ye Chu, Lori M. Lincoln, Gregory E. MacDonald, Noelle L. Anglin, Sudhansu Dash, David J. Bertioli, David Fernández-Baca, Michelle A. Graham, Steven B. Cannon, Ethalinda K.S. Cannon

**Affiliations:** Interdepartmental Genetics and Genomics, Iowa State University, Ames, IA, USA; ORISE Fellow, Corn Insects and Crop Genetics Research Unit, USDA-ARS, Ames, IA, USA; Agronomy Department, Iowa State University, Ames, IA, USA; USDA - Agricultural Research Service, Stillwater, OK, USA; Institute of Plant Breeding, Genetics, and Genomics and Department of Horticulture, University of Georgia, Tifton, GA, USA; USDA - Agricultural Research Service, Corn Insects and Crop Genetics Research Unit, Ames, IA, USA; University of Florida, Gainesville, FL, USA; International Potato Center, Lima, Peru; National Center for Genomic Resources, Santa Fe, NM, USA; Department of Computer Science, Iowa State University, Ames, IA, USA

**Keywords:** peanut, Arachis, genotype, germplasm core collection

## Abstract

Cultivated peanut (*Arachis hypogaea*) is an important oil, food, and feed crop worldwide. The USDA peanut germplasm collection currently contains 8,982 accessions. In the 1990s, 812 accessions were selected as a core collection on the basis of phenotype and country of origin. The present study reports genotyping results for the entire available core collection. Each accession was genotyped with the Arachis_Axiom2 SNP array, yielding 14,430 high-quality, informative SNPs across the collection. Additionally, a subset of 253 accessions was replicated, using between two and five seeds per accession, to assess heterogeneity within these accessions. the genotypic diversity of the core is mostly captured in five genotypic clusters, which have some correspondence with botanical variety and market type. There is little genetic clustering by country of origin, reflecting peanut’s rapid global dispersion in the 18th and 19th centuries. A genetic cluster associated with the *hypogaea/aequatoriana/peruviana* varieties, with accessions coming primarily from Bolivia, Peru, and Ecuador, is consistent with these having been the earliest landraces. The genetics, phenotypic characteristics, and biogeography are all consistent with previous reports of tetraploid peanut originating in Southeast Bolivia. Analysis of the genotype data indicates an early genetic radiation, followed by regional distribution of major genetic classes through South America, and then a global dissemination that retains much of the early genetic diversity in peanut. Comparison of the genotypic data relative to alleles from the diploid progenitors also indicates that subgenome exchanges, both large and small, have been major contributors to the genetic diversity in peanut.

All data is available at the National Ag Library: https://doi.org/10.15482/USDA.ADC/1518508 and at PeanutBase: https://peanutbase.org/data/public/Arachis_hypogaea/mixed.esm.KNWV

## Introduction

Cultivated peanut (*Arachis hypogaea*) was domesticated in central South America by early agriculturalists, following tetraploidization of a hybrid involving the merger of two progenitor diploid species: *A. duranensis* and *A. ipaënsis* (Bertioli et al. 2016). *A hypogaea* has been taxonomically classified into two subspecies, *hypogaea* and *fastigiata*, and several botanical varieties. A period of several thousand years of domestication and diversification in South America led to the establishment and dispersal of several distinct botanical types by the time of Portuguese, Spanish, and Dutch incursion into South America in the 1500s. Establishment of diverse botanical types prior to European contact is evidenced by archaeological records from several locations in South America, including the *hypogaea* and *vulgaris* botanical varieties from regions corresponding with Chile, Argentina, Ecuador, Paraguay, Bolivia, and Brazil (Krapovickas and Vanni 2009); and *peruviana, aequatoriana*, and *hirsuta* varieties from northern South America - now corresponding with Peru, Bolivia, and Ecuador (Krapovickas 1995). Throughout the colonial period (∼1492–1832), peanut cultivation spread quickly around the world. Peanut is now an important source of protein and oil worldwide. In 2017, the 718,570 hectares in the U.S. produced 47,097,498 metric tons; and worldwide, 28 million hectares produced 47 million metric tons (https://www.nass.usda.gov). As a nitrogen-fixing legume, peanut is also important as a rotation crop that restores soil nitrogen.

The USDA peanut germplasm collection provides an essential source of diverse genetic material for breeders. The collection, representing peanut introductions from around the world and most of the ∼80 diploid *Arachis* wild relatives, currently contains 8,982 accessions, which are maintained by the USDA Plant Genetic Resources Conservation Unit in Griffin, GA. As a recent polyploid that experienced a domestication bottleneck, genetic variation across peanut landraces is expected to be low. Peanut is susceptible to a wide range of pathogens, so breeding for disease resistance is of paramount importance. Other traits are important breeding targets, including agronomic traits such as time to maturity and pod-fill, flavor, and nutritional and market traits such as seed size and oil quality.

The U.S. Peanut Core Collection was developed using geographic origin and phenotypic characteristics to select a representative set of accessions from the US collection that span the diversity of cultivated peanut (Holbrook et al. 1993). The development of the Affymetrix SNP array, ‘Axiom_Arachis2’ (Clevenger et al. 2018; Korani et al. 2019) enabled low-cost analysis of this core set through genotyping. The resulting data set will serve multiple purposes: to assess the genetic diversity of the core collection and its population structure; to provide breeders with genotype data for each accession; and to generate data that can be used for trait association (GWAS) analyses. In addition to these expected outcomes, investigation of the phylogenetic and network characteristics of the collection provide information about the historical spread of peanut diversity globally.

The specific objectives of this study were to 1) provide genotype data for each accession, 2) assess genetic diversity of the collection, 3) analyze population structure, 4) estimate the incidence of heterogenous or mixed accessions, and 5) assess relationships between genotypic groups and common traits and phenotypic classes.

## Materials and Methods

### Germplasm material

The U.S. Peanut core collection of 831 accessions was developed in the 1990s. Of these, 44 were unavailable at the time of this study. This project genotyped the 787 accessions which were available (Supplementary File S1) and 14 commercial varieties used in many U.S. breeding programs. These included Tifguard / PI 651853 (Holbrook et al. 2008), Georgia-06G / PI 644220 (Branch 2007b), FloRun 107 / PI 663993 (Tillman and Gorbet 2015), Bailey / PI 659502 (Gorbet and Tillman 2009; Isleib et al. 2011; Tillman and Gorbet 2015), Florida Fancy / PI 654368 / PVP #200800231 (Branch 2007a), Jupiter (Anon. 2000), Tamnut OL 06 / PI 642850 (Baring et al. 2006), OLin / PI 631176 (Simpson et al. 2003), Tamrun OL 11 / PI 665017 (Baring et al. 2013), Red River Runner / PI 665474 (Melouk et al. 2013), NM309-2 (released as NuMex-01) / PI 670460 (Puppala and Tallury 2014, Chamberlin et al. 2015), Florida-07 / PI 652938 (Gorbet and Tillman 2009), Tifguard / PI 651853 (Simpson et al. 2003; Holbrook et al. 2008), and OLé (Chamberlin et al. 2015).

Each accession was grown to maturity to enable seed collection. The accessions which originated from Africa were grown by the Ozias-Akins lab in Tifton, GA. The remaining accessions were grown by the Chamberlin lab in Stillwater, OK. Additionally, we selected 247 accessions for replicate genotyping to test accession purity. These were grown to seedling stage in Ames, IA. Of the 253 accessions, 35 were selected based on information from GRIN-Global (https://www.grin-global.org) and previous knowledge of heterogeneity (Otyama et al. 2019). The remaining 212 were randomly selected to evaluate overall homogeneity of the core collection (Supplementary File S1).

For the replicated genotyping, two seeds were randomly picked from a seed packet of 30 seeds per selected accession. These were then planted in the greenhouse, on a sand bench, or in a growth chamber. Not all selected samples germinated (even after replanting), which limited the number of samples available for replicate genotyping for some accessions. Of the 247 accessions; 197 accessions were genotyped twice, 33 were genotyped three times,16 accessions had four samples and one had five samples genotyped. In total, 1145 samples were available for genotyping.

### DNA extraction and genotyping

For all accessions, whether grown to maturity or to seedling stage, leaf tissue was sampled between 2 and 4 weeks after germination and immediately frozen in liquid nitrogen. DNA was extracted using Qiagen (Germantown, MD) DNeasy 96 Plant Kits (#69181) and 3 mm Tungsten Carbide Beads (#69997) as recommended by the manufacturer. Initial concentration and purity of 12 DNA samples/plate was estimated using a Thermo Fisher Scientific® NanoDrop ND-1000 Spectrophotometer (Thermo Fisher Scientific®, Waltham, MA, USA). Concentrations ranged from 26 to 75 ng/ul, with an average 43 ng/ul. A260/A280 ratios ranged from 1.882 to 1.984, with an average ratio of 1.931. A260/A230 ranged from 1.84 to 2.681, with an average ratio of 2.206. Samples were then shipped to Thermo Fisher ® for additional quality control and genotyping. DNA concentration and quality for all samples was confirmed using a ‘PicoGreen’ assay. Average DNA concentration was about 47 ng/µL for 926 high-quality samples. The remaining 219 samples had a concentration of 15 ng/ul and were considered of sufficient quality and quantity for genotyping. Samples were then genotyped using the 48k Thermo Fisher ® ‘Axiom_arachis2’ SNP array. Of the 1,145 samples, 25 replicate samples were not successfully genotyped.

Raw SNP intensities from Affymetrix were analyzed using the ‘Best Practice Workflow’ available in the Axiom Analysis Suite. A total of 47,837 SNPs was obtained, of which 14,430 were categorized as ‘Poly High Resolution’, 15,528 were ‘Mono High Resolution’, 11,008 were ‘No Minor Homozygote’, and the remaining 6,871 were of low-quality. Poly High Resolution SNPs were processed into a standard VCF format (Supplementary Files S2 and S3) using custom bash scripts for downstream analyses (https://github.com/cannongroup/peanut_core_collection_genotyping).

### Diversity, phylogenetic, and network analysis

Several aspects of diversity analysis were carried out on variant data in FASTA format - i.e. with SNP variants represented as DNA bases, positioned in the genomic order of the loci. A FASTA-format sequence representation of the ‘Axiom_Arachis2’ SNP array variant data was generated by converting genotype calls in the array to DNA base calls from the Axiom_Arachis2 VCF file generated by ThermoFisher, using custom shell scripts that converted AA/BB calls to A, T, C, G, or “-” (scripts are available at https://github.com/cannongroup/peanut_core_collection_genotyping). The matrix contains 14,430 high-confidence SNPs, for 1,120 samples. Relative positions of the SNPs were also determined from the consensus genomic locations from five *Arachis* genome assemblies, as described below. This sequence representation is available as Supplementary Files S4 and S5.

Base-calls were also derived computationally for four sequenced *Arachis* genomes: *A. duranensis, A. ipaënsis* (Bertioli et al. 2016) and *A. hypogaea* varieties Tifrunner (Bertioli et al. 2019), Shitouqi (Zhuang et al. 2019), and Fuhuasheng (Chen et al. 2019). Base-calls from the genomic sequences were made by aligning flanking sequences plus the variant base, using two sequences per variant per locus, to the respective genome, using blastn (Altschul et al. 1990). Per-locus SNPs were called when the flanking+variant sequence matched at 100%, over at least 65 of 71 bases, to only one location in the genome (i.e. full-length alignments were not required, but perfect match was required within the alignment).

The genome-derived SNPs were added to a version of the sequence variant-call file (Supplementary Files S4 and S5) with the *A. duranensis* and *A. ipaënsis* calls combined into one “synthetic-tetraploid” accession. Base calls that were absent in that accession were removed from the merged file, giving an alignment 10,278 bases wide, by 1,123 samples (after removal of PI493562_1, which appears to have had a label tracking error). Approximate genomic locations of SNPs were determined as: the location in the respective diploid chromosomes were present; otherwise, the location in Tifrunner; otherwise in Shitouqi; otherwise the location in Fuhuasheng, as shown in Supplementary File S6. Two reduced alignments were also generated (Supplementary Files S7 and S8), consisting of representative “centroid” sequences from clusters at the 98% and 99% identity levels, using the cluster_fast method in the vsearch suite, version 2.4.3 (Rognes et al. 2016).

The phylogenetic tree in Figure 1 and Supplementary File S9 was calculated using FastTreeMP, version 2.1.8 (Price et al. 2010), with default parameters. The network diagram in Figure 2 was calculated on the 99%-identity centroid alignment, using the Neighbor-Net algorithm in the SplitsTree package, version 4.15.1 (Huson and Bryant 2006).

**Figure 1.**
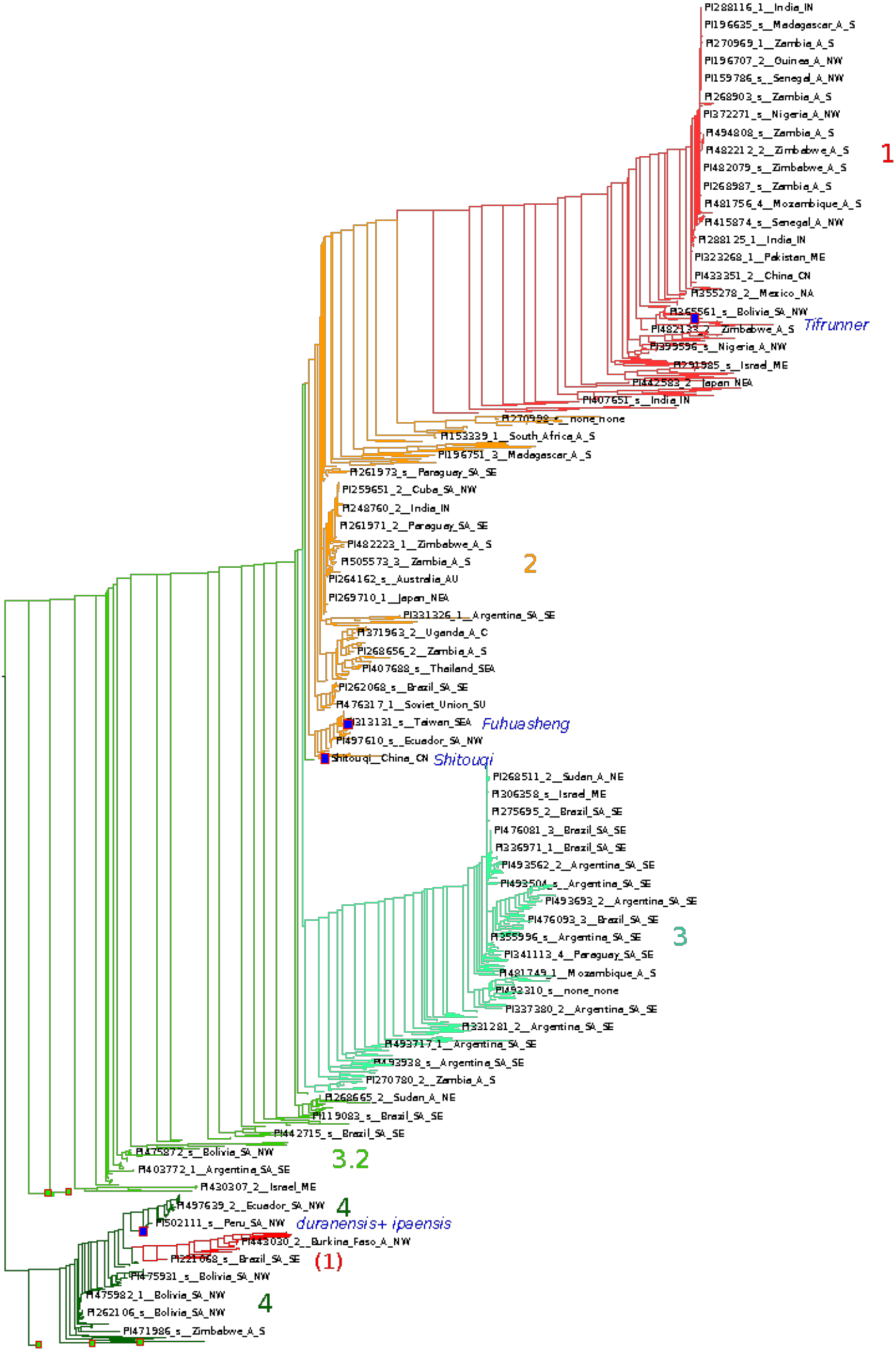
Phylogenetic tree for 1122 samples from 791 accessions of the U.S. peanut core collection. For reference, five clades have been assigned (1-4 and a transitional group, 3.2). These clade designations are also used in the network plot (Figure 2) and in the PCA analysis (Figure 4)

**Figure 2.**
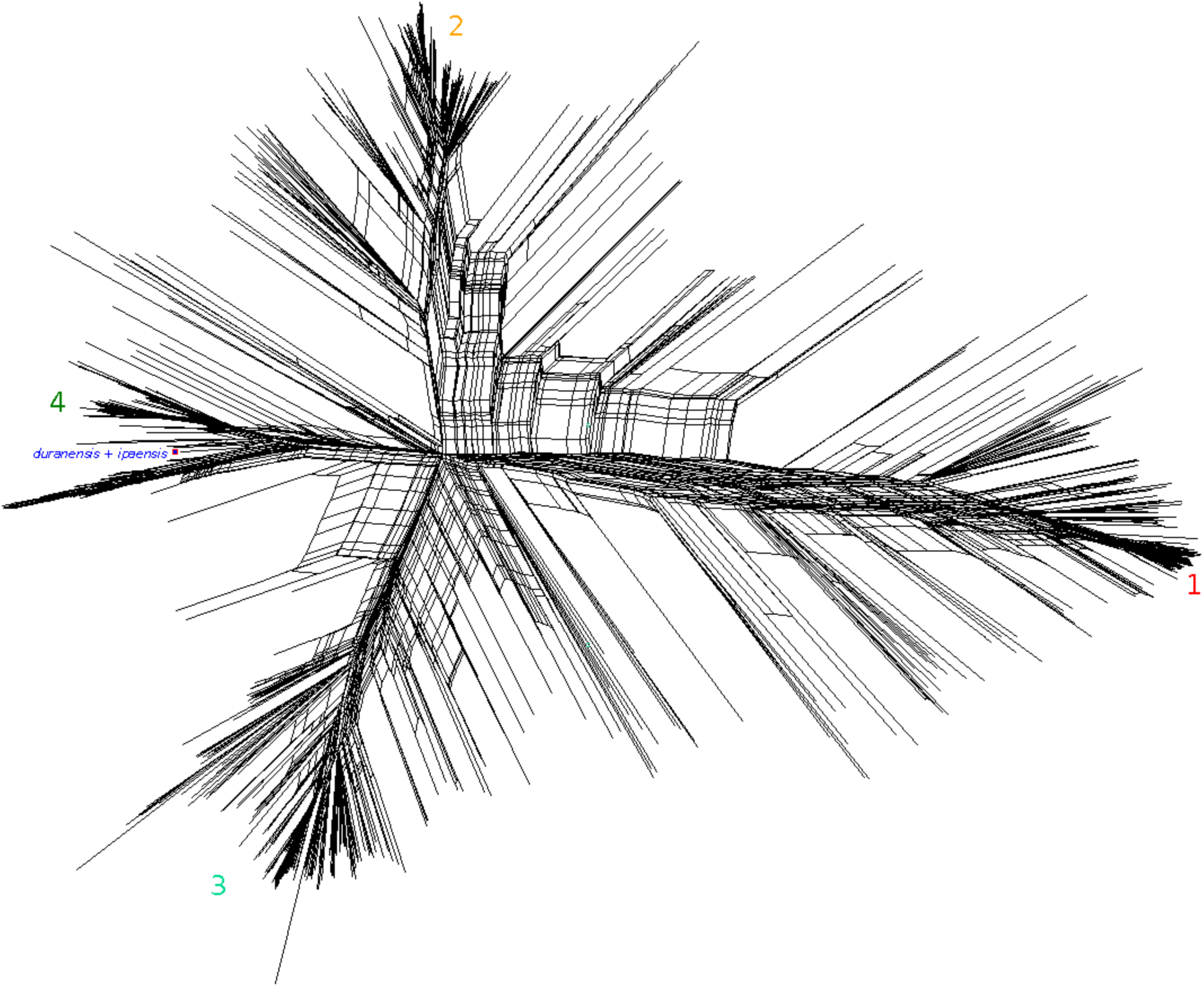
Phylogenetic network of 1122 samples from 791 accessions of the U.S. peanut core collection. Network analysis was performed in SplitsTree using the Neighbornet algorithm with default settings. Accessions are ordered as in the phylogenetic clade analysis with four main clades shown in the figure.

### Replicate analysis

To assess the genetic similarity among multiple samples from an accession, a list of all possible pairs of replicates per accession was calculated, giving “N choose 2” combinations for an accession with N samples: 3 combinations for an accession with 3 samples; 6 combinations for an accession with 4 samples, etc. For each possible combination, the sequence identity was calculated between the sequence pairs (using blastn), and then scored as “similar” if >= 98% identity and “dissimilar” otherwise. These results are shown in the “rep analysis” worksheet of Supplementary File S1.

### Structure and Principal Component Analysis (PCA)

To define subpopulations based on genomic sequences, a structure analysis and PCA was performed on high confidence Axiom_Arachis2 SNP array variant data. Structure analysis was performed using a Bayesian inference algorithm implemented in fastStructure (Raj et al. 2014).

The fastSTRUCTURE resulted in five clusters (K=5) which are shown in Figure 3 and Supplementary Files SF10 and SF11. All 13,410 SNP sequences were used for a representative set of 518 “unique” accessions, selected based on sequence identity at 98%. Clusters and group membership were determined for arbitrary groups ranging from K 1 to 10 with settings: *--prior = logistic, --cv = 0, --tol = 10e-6*, default otherwise. Structure was visualized as proportionally colored bar plots representing global ancestry estimates (Q values) using an R package, Pophelper version 2.3 (Francis 2017).

**Figure 3.**
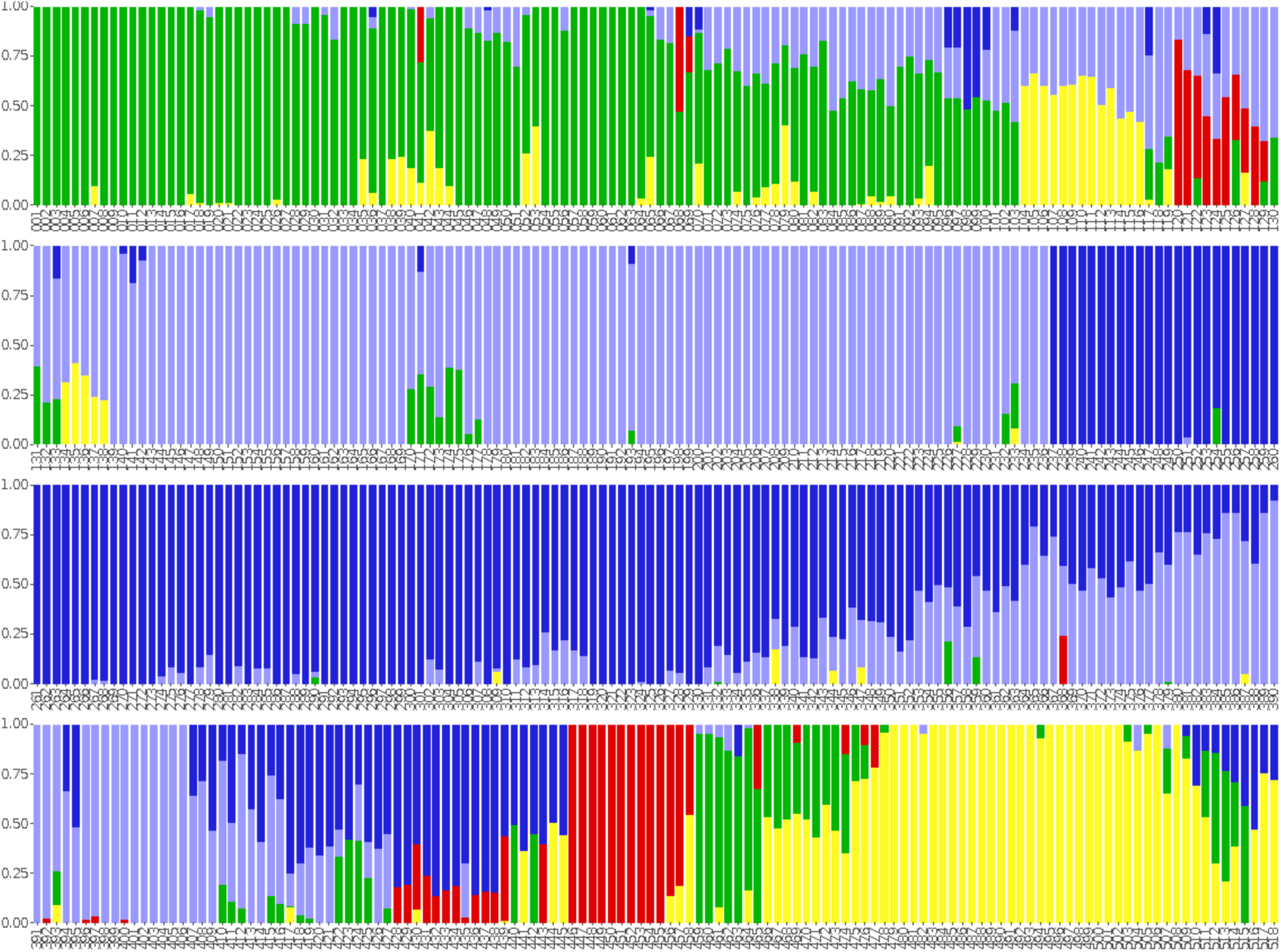
Genetic structure of 518 samples selected as representatives at >= 98% sequence identity. Accessions are grouped into five clusters represented by distinct colors. The X-axis represents accessions ordered according to their positions in the phylogenetic tree analysis. The Y-axis represents proportions of cluster assignment based on Q values from fastStructure analysis.

To avoid the strong influence of SNP clusters in principal component analysis (PCA) and relatedness analysis, only SNPs in approximate linkage equilibrium with each other (r^2^ = 0.2) were used. The R package, SNPRelate (Zheng et al. 2012), was used for LD pruning on 1120 samples. snpgdsLDpruning in the SNPRelate package, was used to recursively remove biallelic SNPs in LD within a sliding window of 1Mb. LD threshold was specified at r^2^ = 0.2. Monomorphic SNPs were also removed along with uncommon SNPs filtered at MAF < 5% leaving a final set of 2,063 markers in approximate linkage equilibrium with each other. PCA was performed using snpgdsPCA from the SNPRelate package at default settings and plotted using ggplot2 for defined groups. PCA results are shown in Figure 4 and Supplementary File S12. Groups were defined according to: whether or not they flowered on the main stem, their botanical variety defined in GRIN-Global, agronomic type (market group), growth form, pod type, and country from which seed was originally collected.

**Figure 4.**
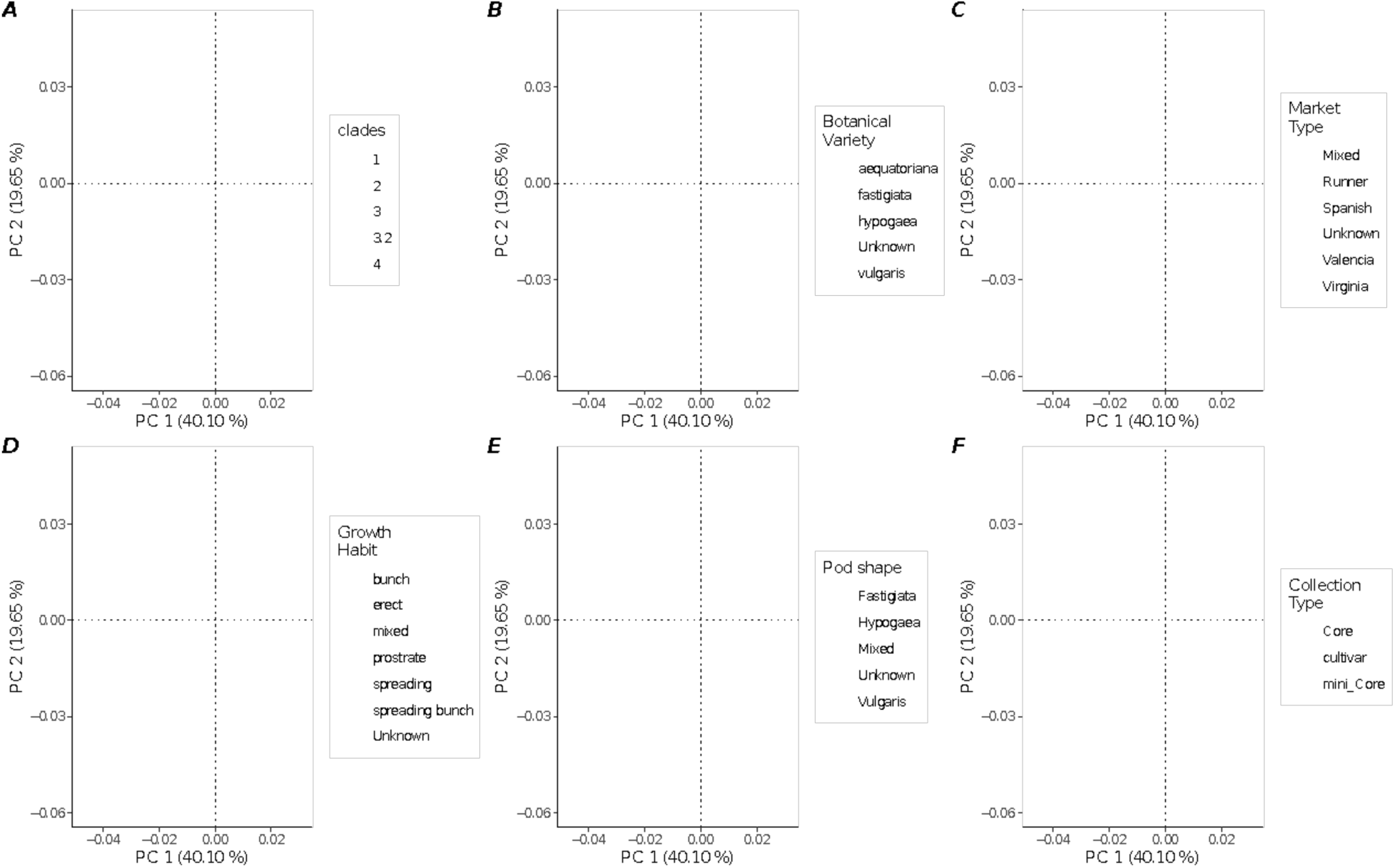
Principal Component Analysis of 1120 samples based on 2063 unlinked SNP markers. The X-axis represents PC 1 and the Y-axis represents PC 2. Samples are colored and grouped according to: A. clade membership as defined in the phylogenetic and network analyses, B. botanical varieties, C. market type, D. growth Habit, E. pod shape, and F. collection type

### Population differentiation analysis

To evaluate differentiation between and among accession groups, we calculated F_ST_ for selected accession groups defined as above under the Structure and PCA methods section. Results are shown in Figure 5A-F. SNPs were first pruned to reduce SNPs in strong LD with one another, as described above. The F_ST_ analysis was performed using the R package Hierfstat, (Goudet 2005) at default settings. Pairwise F_STs_ were calculated using pairwise.WCfst according to (Weir and Cockerham 1984). A heatmap of pairwise F_STs_ was plotted using ggcorrplot (Kassambara 2016), for defined groups.

**Figure 5.**
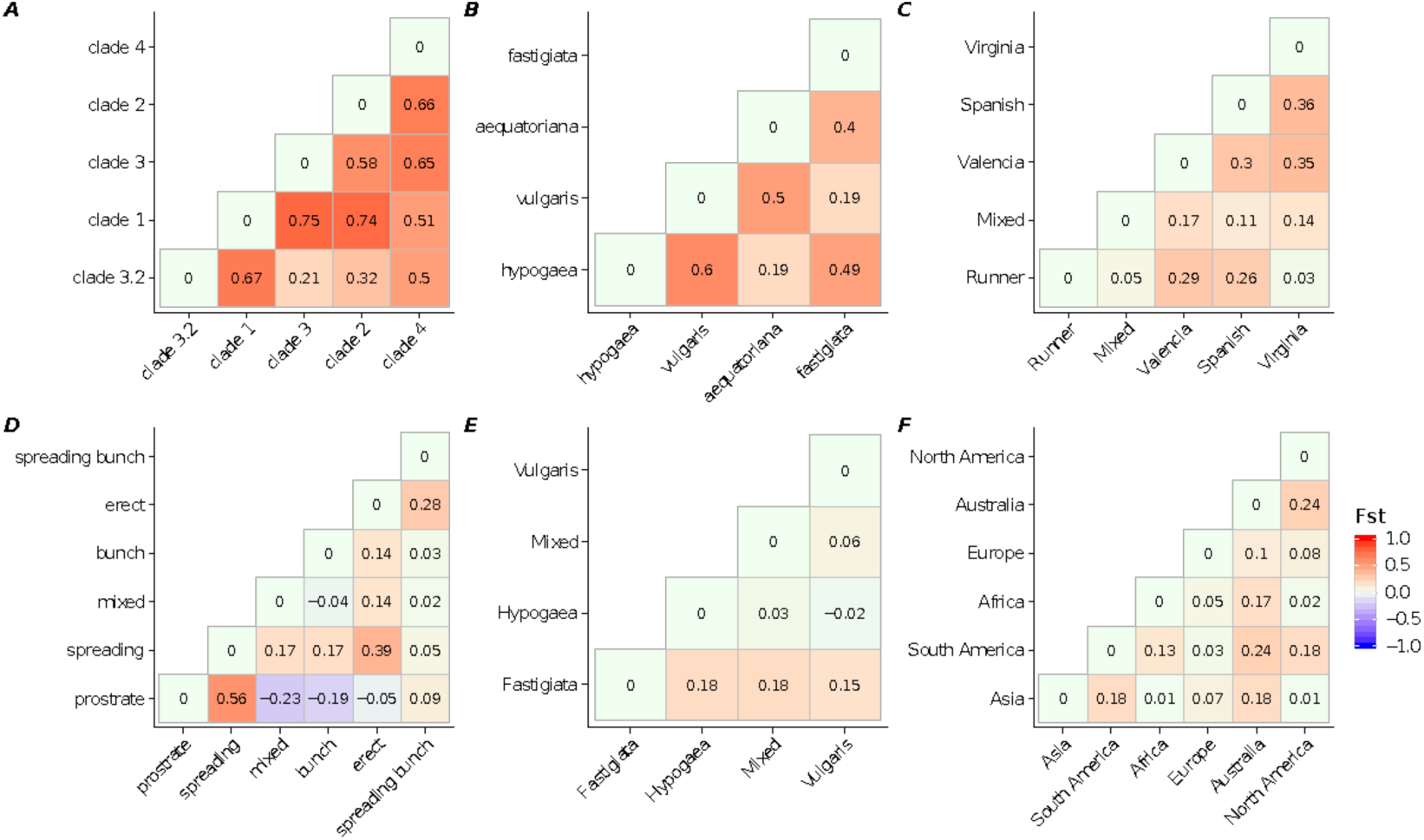
Plots of F_ST_ (fixation index) values among genetic groupings, to determine stratification in the core collection. Cluster identities are as shown in the phylogenetic and PCA analyses. The pairwise population differentiation (F_ST_ index) was calculated using Hierfstat for a set of unlinked markers and plotted as heatmaps. Accessions were classified into groups of: A. clade membership as defined in the phylogenetic and network analyses, B. botanical varieties, C. market type, D. growth Habit, E. pod shape, and F. continent of seed origin.

### Geographical distribution

A plot of the geographical distribution of peanut accessions by clade (Figure 6) was generated using the germplasm Geographical Information System (GIS) utility at PeanutBase.org (Dash et al. 2016), with the “add your data” tool. To display the five germplasm categories identified in Figures 1 and 2, we used the following column labels, which are interpreted by the GIS tool: accession_id, trait_observation_value, trait_descriptor, taxon, trait_is_nominal.

**Figure 6.**
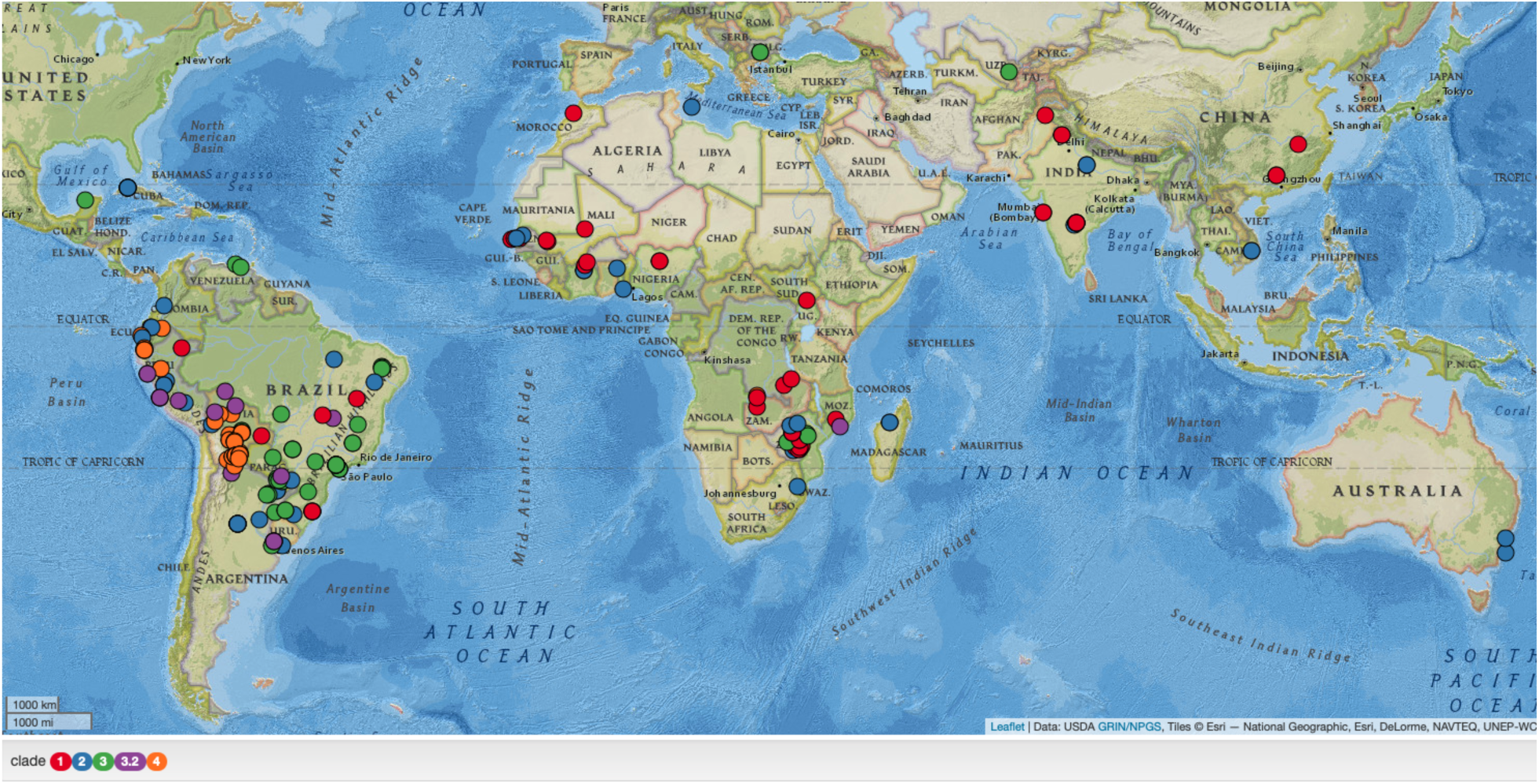
Geographic origin of genotyped accessions. Colors indicate clades in Figure 1 (colors and clade correspondences are shown in the legend in the lower left in the figure). Figure was generated using the Germplasm GIS tool at peanutbase.org.

### Analysis of subgenome invasions

To track possible instances of subgenome interactions, 16 accessions were selected from across the clades identified in Figures 1 and 2 and alleles were examined relative to those identified in the diploid accessions (Supplementary Files SF6 and SF13). Alleles for each accession were then marked as being the same as the A-genome allele and not the B-genome allele (A-like), or same as the B-genome allele and not the A-genome allele (B-like), or other conditions (invariant in the diploids, different from both diploids, or missing in one or more of the tetraploid or diploid accessions). The results are shown in Figure 7, with red indicating identity with the respective subgenome (A-like for chromosomes 1-10, and B-like for chromosomes 11-20).

**Figure 7.**
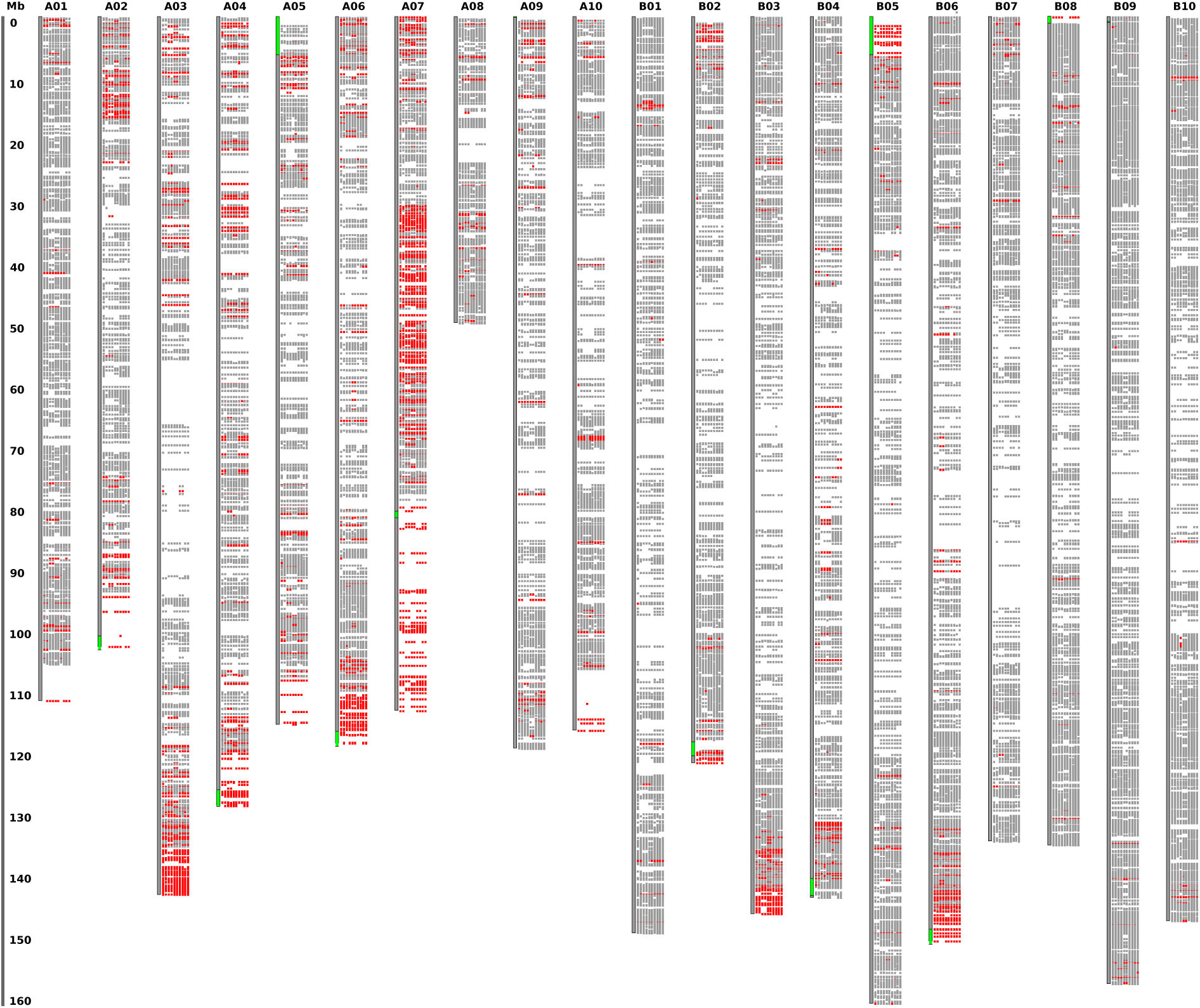
Plot of inferred subgenome origins. Each colored region (gray or red) indicates data at a SNP location. At each position, values are shown for 16 diverse accessions. In chromosomes A01-A02 (left half), red indicates that alleles are the same as the B-genome assembly (A. ipaensis) and different than the A-genome assembly (A duranensis), at the respective locations (determined by perfect correspondence of flanking sequence). In chromosomes B01-B10 (right half), red indicates that alleles are the same as the A-genome assembly (A. duranensis) and different from the B-genome assembly (A ipaensis). Green marks on the chromosome backbones (e.g. tops of A05 and B05) show the locations of large-scale subgenome invasion, observed in the Tifrunner genome assembly (Bertioli et al., 2019).

### Data Availability

All data is available at the National Ag Library: https://doi.org/10.15482/USDA.ADC/1518508

and at PeanutBase: https://peanutbase.org/data/public/Arachis_hypogaea/mixed.esm.KNWV.

Scripts used in data analysis are available at GitHub: https://github.com/cannongroup/peanut_core_collection_genotyping.

**File S1** (tables) [SF01_peanut_core_v14.xlsx] contains the main descriptive information about the genotyped accessions, including: information about replicate similarity; phylogenetic clades, geographic origin, and phenotype; and summaries of phenotypic and country information relative to clade assignments.

**File S2** (text file) [SF02_SNPs_whole_Axiom_Arachis2.txt] has the original genotype calls for the Axiom array (for poly-high resolution SNPs).

File S3 (text file) [SF03_SNPs_whole_Axiom_Arachis3.vcf] has the Axiom array genotype calls, in VCF format.

**File S4** (text file) [SF04_SNPs_w_4_genomes.tsv] has the predominant DNA variants at each SNP location, for all accessions, including variants inferred from four available genome assemblies: *A. duranensis* and *A. ipaensis* together, and *A. hypogaea* accessions Tifrunner, Shitouqi, and Fuhuasheng. The format is in a simple tab-separated table, with 14431 columns (SNP positions).

**File S5** (text file) [SF05_SNPs_w_4_gnm_mrgd.fas] the same SNP as in S4 above, but in fasta format. SNP locations without DNA assignments for *A. duranensis* and *A. ipaensis* have been removed, giving an alignment of 10278 bases.

**File S6** (tables) [SF06_chip_and_genome_samples_v04.xlsx] has DNA base-calls for 16 selected, diverse accessions, with comparisons to the variants observed in the *A. duranensis* and *A. ipaensis* genomes, and inferences regarding the likely progenitor for the DNA, i.e. A-genome (*A. duranensis*) or B-genome (*A. ipaensis*).

**Files S7 and S8** (text files) [SF07_SNPs_w_4_gnm_mrgd_cen98.fas and SF08_SNPs_w_4_gnm_mrgd_cen99.fas] are reduced fasta alignments (relative to the complete alignment file, S5). File S7 has the centroid representatives at 98% identity, and S8 has centroid representatives at 99% identity. These files have 518 and 680 sequences, respectively.

**File S9** (text file) [SF09_SNPs_w_4_gnm_mrgd_rt3.nh.txt] is the phylogenetic tree (Newick format) calculated from the alignment in S5, and corresponding with the phylogenetic tree shown in Figure 1.

**File S10** (figure) [SF10_K5_membership.pdf] shows the proportion of accessions assigned to clusters 1-5 in a Structure analysis (Figure 3), for K=5 clusters.

**File S11** (tables) [SF11_K5_cluster_assignment.xlsx] gives the proportional assignments of each cluster to all accessions (relative to the Structure diagram shown in Figure 3).

**File S12** (figure) [SF12_pca_34.pdf] Principal Component Analysis of 1120 samples based on 2063 unlinked SNP markers. The X-axis represents PC 3 and the Y-axis represents PC 4. Samples are colored and grouped according to: A. clade membership as defined in the phylogenetic and network analyses, B. botanical varieties, C. market type, D. growth habit, E. pod shape, and F. collection (core, mini core, cultivar).

**File S13** (tables) [SF13_chip_and_genome_GFFs.xlsx] Inferred subgenome origins of SNPs relative to the A-genome and B-genome progenitors (*A. duranensis* and *A. ipaensis*). This data is in GFF format, derived from S6, and used as the basis for the plots in Figure 7 (showing regions of possible subgenome invasions).

**File S14** (figure) [SF14_PI497426_pods.jpg] Pods from accession PI 497426 (clade 4), illustrating the distinctive reticulation pattern seen in some accessions in this clade.

**File S15** (figure) [SF15_Sipan_neclkace_Donnan_Einstein.jpg] Picture of necklace of peanuts, sculpted in gold and silver, from the Moche-era tomb at Sipán (ca. AD 250) in coastal Peru. Photograph by Susan Einstein, courtesy of Christopher Donnan.

## Results and Discussion

### Replicate analysis

For the 253 accessions with replicates, a maximum of 428 pairings from same-accession groupings were expected. For example, an accession with one replicate (A and B) has one expected pairing (A-B), while an accession with two replicates (A,B,C) has three expected pairings (A-B, A-C, B-C), and an accession with three replicates has six expected pairings. A missed pairing means that one or more samples for an accession are genetic outliers, and that the accession is not homogeneous. Accessions chosen for replicate genotyping included 35 accessions noted in GRIN-Global as being potentially mixed or in which the seeds appeared to be visibly heterogeneous. Additionally, replicate genotyping was carried out for 218 accessions selected at random from the core.

Of the 428 expected pairings among replicates (with >70% sequence identity across all SNP locations), 368 pairings were observed (86%). The observed pairings had an average identity of 94.4% and a median of 98.7%. The 60 instances of a sample that did not match to a replicate for that accession occurred among 42 accessions, meaning that some accessions had more than one “missing” match for a replicate.

Of the 35 accessions selected as “probably mixed” based on seed color or other notes in GRIN records, most (77%) were indeed mixed genotypically only eight of these accessions had all of the replicates close to identical (>=98%) across all replicates. For the others (27/35), at least one sample per accession was not like the others at the 98% identity threshold.

Of the 236 accessions selected at random for replicate genotyping, most (56%) accessions were NOT mixed genotypically: in 123 of these accessions, all replicates were close to identical (>=98%) across all replicates. Nevertheless, the high rate of apparent genotypic heterogeneity in accessions suggests that the core collection will require further subdivisions or selections to generate material that is well suited for analyses such as QTL and GWAS.

### Diversity analysis: phylogenetic analysis

The core collection contains considerable phenotypic diversity, but also displays high genotypic similarity among many accessions, as apparent in Figure 1, where many accessions are near-identical in the phylogeny. The 1,122 samples (791 accessions) in this study fall into 671 clusters at an identity threshold of 99% (Supplementary File S1, worksheet “clusters”). The largest clusters at 99% identity have 139, 49, 27, and 25 samples (112, 42, 21, and 22 distinct accessions), and the cluster sizes fall progressively to the singletons, of which there are 560. The existence of large clusters of nearly identical accessions suggests that diversity in the core could be represented by a smaller number of accessions (671, specifically, if 99% identity were used as the identity cutoff).

The phylogenetic tree of accession diversity shows four primary clades of accessions, numbered 1-4 in Figure 1, with an intermediate group (3.2) also indicated. These clade numbers are also used in the network diagram Figure 2. Although some accessions occur on early branches in these clades (rather than nested tightly in terminal clusters), the clades are nevertheless mostly distinct in both the phylogeny and the network plot. The clade designations also generally correspond with the Structure plot at cluster-number K=5 Figure 3. The Structure plot is ordered by the tree order from Figure 1.

A top-level summary of the cluster- and trait-correspondences demonstrates that most accessions, including all named cultivars, fall into three large clades (1, 2, and 3), but those clades don’t correspond cleanly with typical peanut classifications (e.g. growth habit, botanical variety, market type, or pod type). Traits categories are shown superimposed on the clades, in the PCA plots in Figure 4. A smaller clade (4) does correspond with these typical classification traits (Figures 1 and 4). Clade 4 has exclusively erect growth habit, with pod-types of hypogaea, valencia, or mixed pods, but frequently having strong, linear reticulation, and including the *aequatoriana* botanical variety of subspecies *fastigiata*, as exemplified by PI 497426 from this clade (Supplementary File S14).

For each cluster, counts and proportions of phenotypic characters and collection region are given in Table 1. The clusters have some correspondence with growth-habit traits and with countries of seed origin, as described below. (In this section, all counts are given per accession rather than per sample, as some accessions were genotyped multiple times).

**Table 1.**
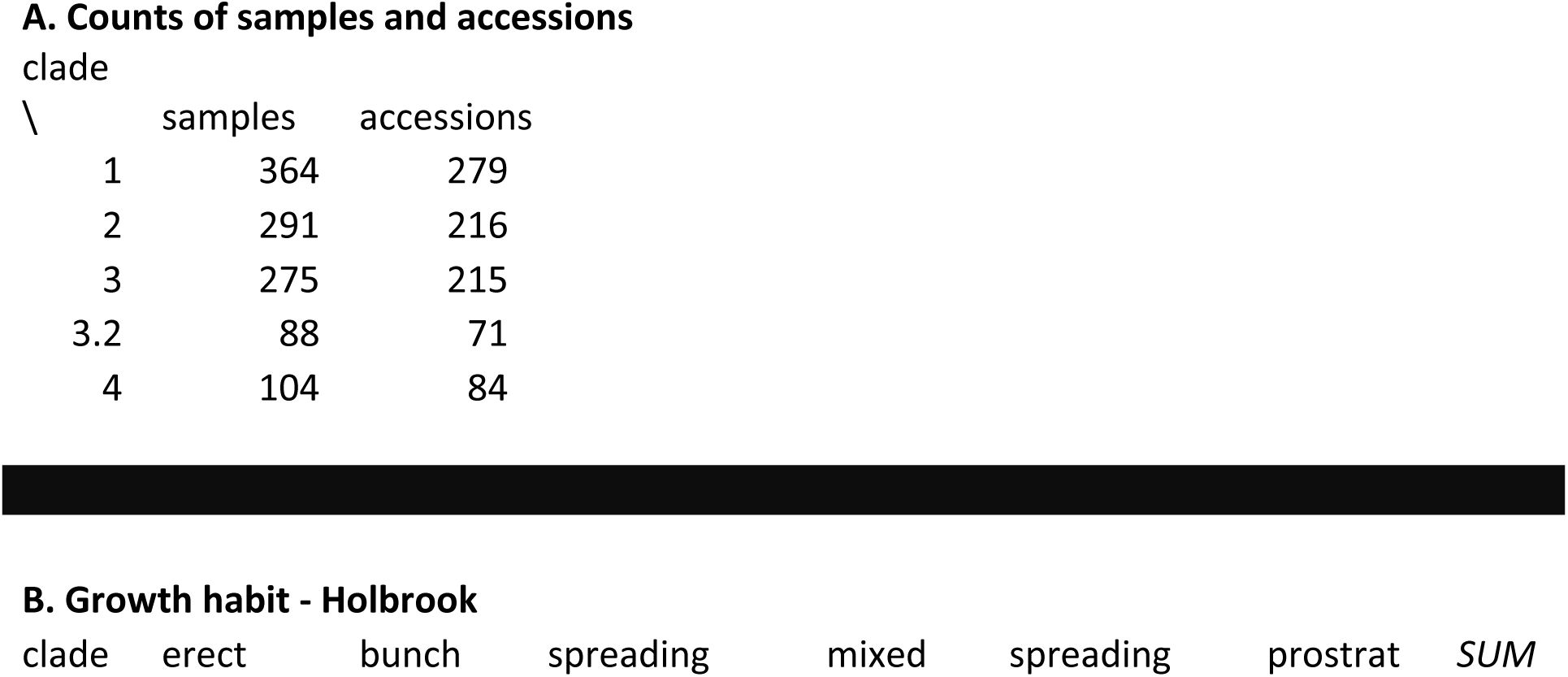
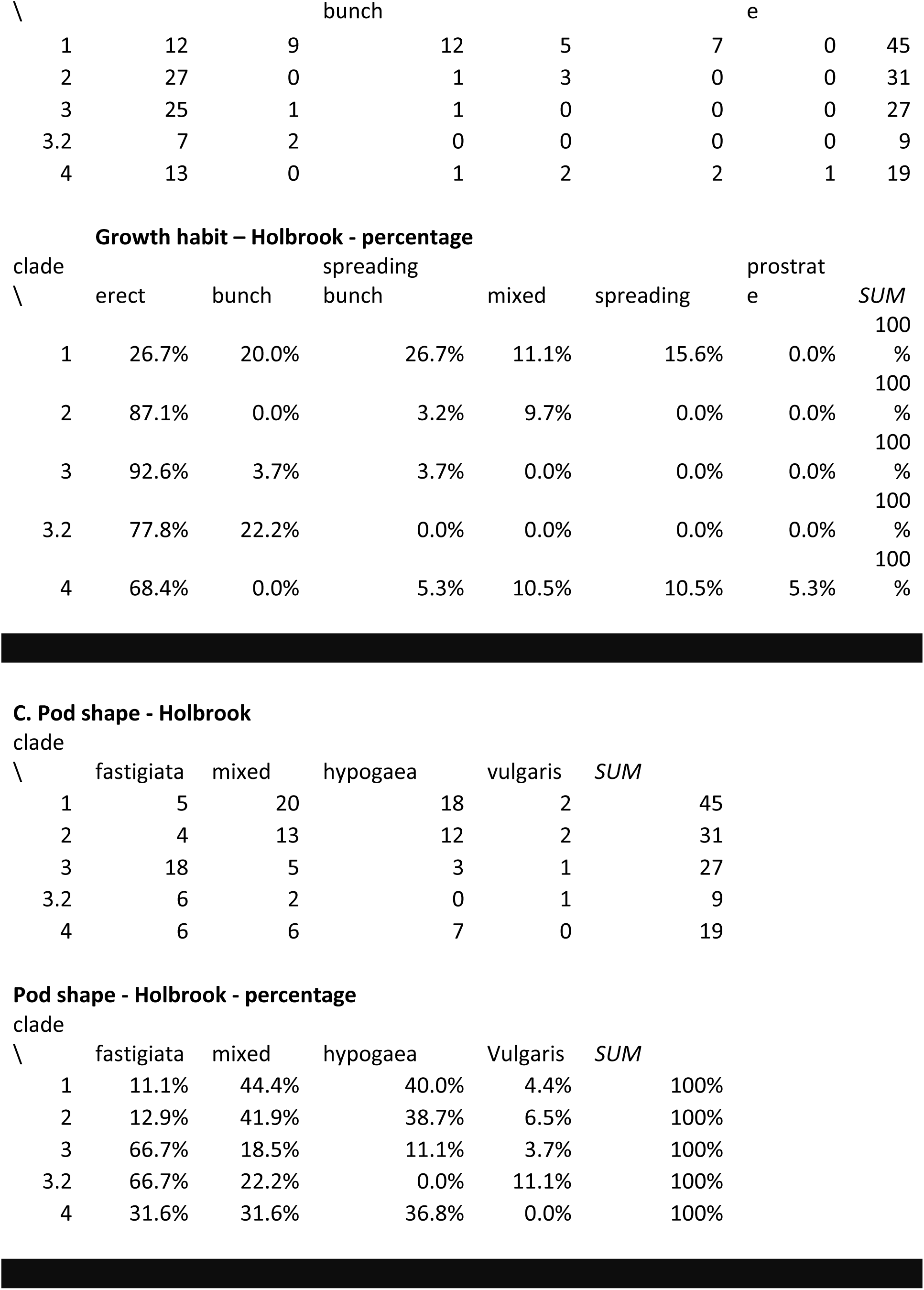
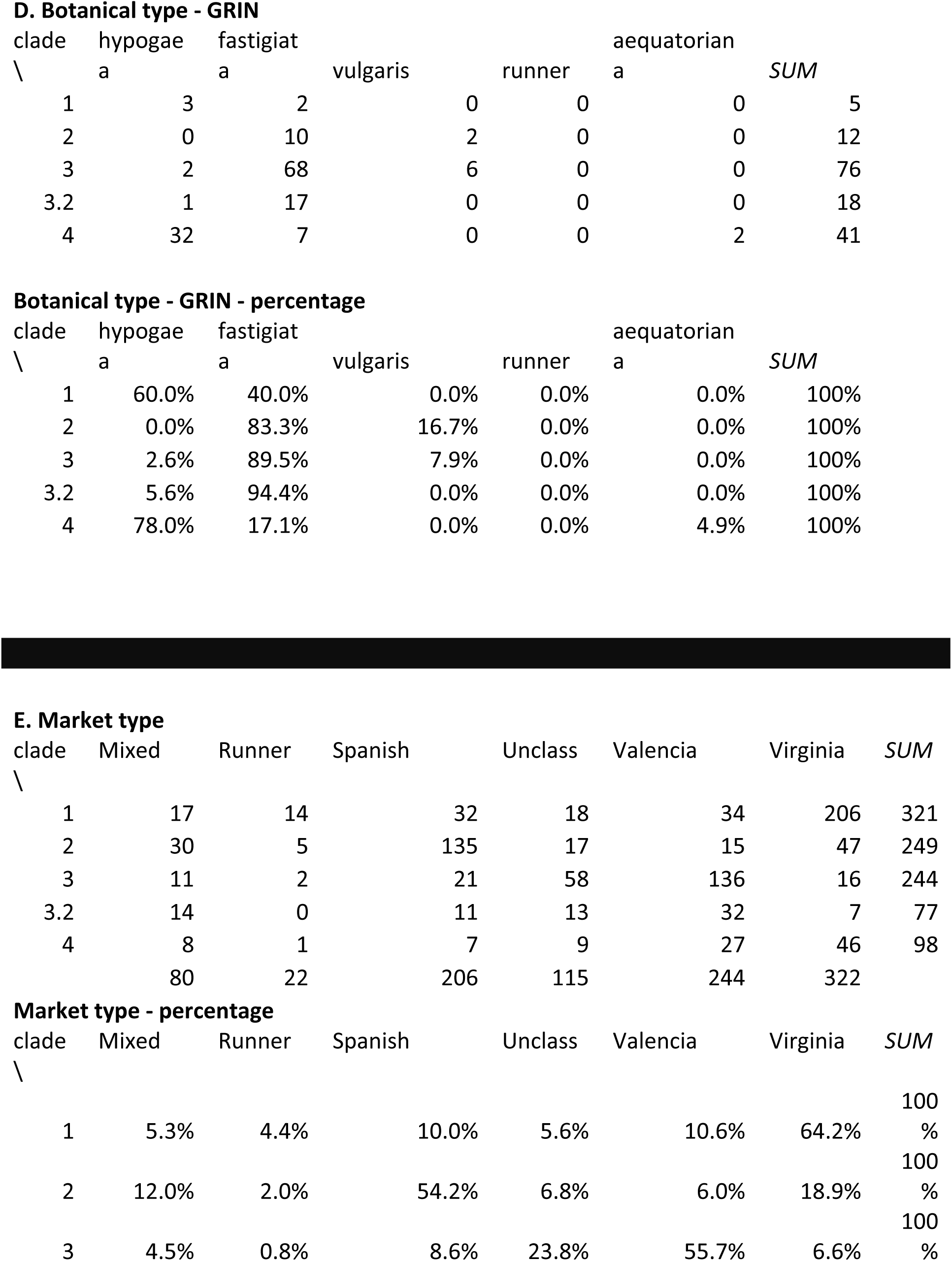
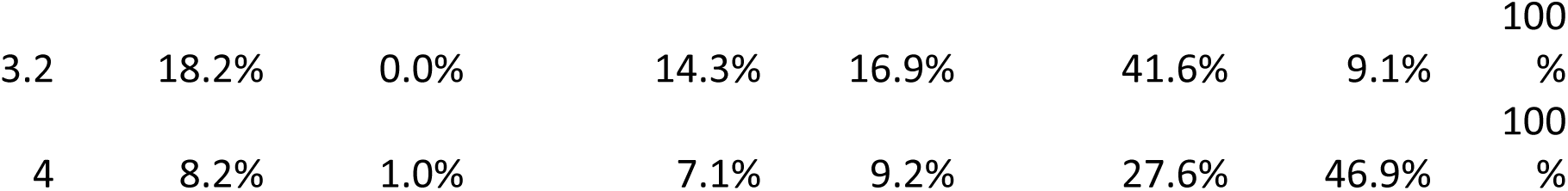
Counts of genetically unique samples, relative to phenotypic traits. Unique samples are listed in Supplementary File S1, worksheet “uniques”. Table 1A: counts of samples and accessions per clade (relative to clades identified in Figure 1). Tables B-E: counts of unique accessions per clade and per trait; trait classes as identified in table subheadings. Traits are per Holbrook et al. (1993) and the Germplasm Resources Information Network (GRIN), as indicated.

<u>Clade 4</u> (Figures 1 and 4; at the bottom in Figure 1; 104 samples, 84 accessions) is the most distinctive and consistent phenotypically: most accessions (68.4%) have upright growth habit, per Holbrook’s phenotype evaluations (Holbrook and Dong 2005). The pod type is more varied, with accessions scored as *hypogaea, fastigiata*, or mixed (36.8, 31.6%, 31.6%)(Holbrook and Dong 2005). Growth type was scored as *fastigiata* for seven accessions and two as *aequatoriana*. The *aequatoriana* type is a botanical variety of the subspecies *fastigiata* (Krapovickas et al. 2007). pod images from GRIN-Global for this clade show pods frequently having strong reticulation and widely-spaced veins running the length of the pod (Supplementary Figure S14) - which is of interest as these characteristics are seen in pre-colonial archaeological finds in Peru, Chile, and Argentina (Supplementary Figure S15). Most of the cluster 4 accessions originate from west-central South America (Figure 6), primarily from Bolivia, Peru, Ecuador, and Argentina (38, 17, and 9, and 8 accessions, respectively). Interestingly, the inferred genotype for *A. duranensis* and *A. ipaensis* (consisting of alleles at loci corresponding with the marker flanking sequences from the SNP array) also falls solidly within cluster 4, with 100% bootstrap support on several subtending branches in this clade.

<u>Clade 3.2</u> (Figures 1 and 4; second from bottom in Figure 1; 88 samples, 71 accessions) shows general phenotypic consistency: most accessions have the *fastigiata* botanical variety, upright growth habit, and *fastigiata* pod type (94.4%, 77.8%, and 66.7%, respectively). This is a transitional clade, with similarities to both Clades 2 and 3.

<u>Clade 3</u> (Figures 1 and 4; third from bottom in Figure 1; 275 samples, 215 accessions) shows general phenotypic consistency: most accessions have the *fastigiata* botanical variety, upright growth habit, and *fastigiata* pod type (92.6%, 66.7%, 89.5%, respectively). Both characteristics distinguish this group from Cluster 4. The most frequent South American accession origins for Cluster 3 are Bolivia, Argentina, and Brazil (40, 5, 5, respectively), with one each from Peru and Ecuador. The most frequent non-South American countries for cluster 3 are Zambia, Nigeria, and Zimbabwe (12, 6, and 6, respectively).

<u>Clade 2</u> (Figures 1 and 4; second from top in Figure 1; 291 samples, 216 accessions). In this clade, most accessions have the *fastigiata* botanical variety, *fastigiata* growth habit, and *fastigiata* pod type (83.3%, 87.1%, 83.3%, respectively). The Clade 2 accessions also have the widest geographic spread. also cosmopolitan in terms of country of origin. The most frequent South American countries for these accessions are Brazil, Argentina, Cuba, and Uruguay (10, 9, 6, 5, 5, respectively). Non-South American countries are the predominant sources for these accessions, however; Zambia, Zimbabwe, India, and Sudan are the most frequent sources (34, 17, 13, 13, 13, respectively). Because the highest-frequency countries of origin are Brazil in South America and Zambia, Zimbabwe and Sudan in Africa suggests early movement of this germplasm through the slave and other colonial trade.

<u>Clade 1</u> (Figures 1 and 4; top in Figure 1; 364 samples, 279 accessions). In this clade, most accessions are classified as the *hypogaea* botanical variety and “mixed” or *hypogaea* pod shape (60.0%, 44.4%, 40.0%). Growth type varies widely, divided fairly evenly between erect, bunch, spreading-bunch, mixed, and prostrate). The most frequent market type is Virginia (64.2%). As with Cluster 2, the geographical spread is highly cosmopolitan (Figure 6), with the largest numbers coming from Zambia, Israel, India, Nigeria, and China (40, 37, 29, 27, 26, respectively).

### The geographic distribution of genotypes

All parts of the phylogenetic tree are dominated by accessions from South America, but all clades also have interspersed accessions from many parts of the world Table 2. This pattern of broad geographical dispersal, with heavy representation in South America, confirms that peanut had fully diversified into modern cultivar types prior to dispersal through colonial shipping and trade. Influence of the slave and spice trade is suggested by adjacent appearance in the phylogenetic tree of widespread geographical locations. for example, accessions from Portugal are interspersed among accessions from countries in west Africa, south Asia, and the Caribbean and eastern South America (in Clades 1, 2, 3, and 3.2) or Spain and countries in Africa, the Middle East, and Asia (middle of Clade 1).

**Table 2.**
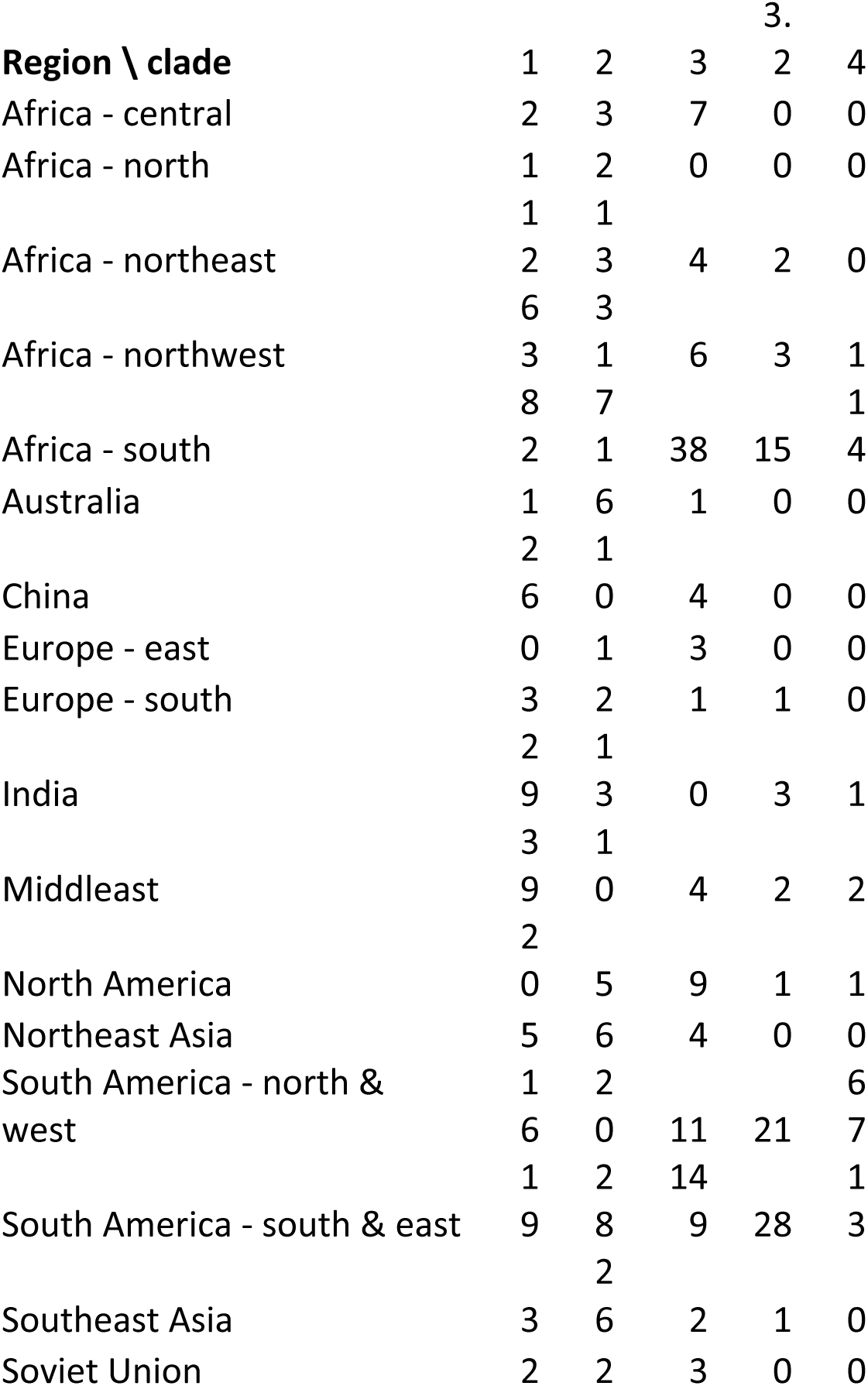
Counts of genetically unique samples, relative to geographic regions. Unique samples and countries and regions are listed in Supplementary File S1, worksheet “uniques.” Detailed counts (per country) are given in Supplementary File S1, worksheet “clade summary.” Columns labeled 1-4 indicate clades, as identified in Figure 1, and listed in File S1, worksheet “uniques.”

Clade 4 is much less mixed geographically, coming predominantly from central and western South America (Figure 6). Peanut’s geographic origin (through the initial instance of tetraploidy) has been convincingly established as having occurred in (Bertioli et al. 2016; Bertioli et al. 2019)southeastern Bolivia/northwestern Argentina (Bertioli et al. 2016; Bertioli et al. 2019). It is therefore noteworthy that the combined diploid progenitors (*A. duranensis* and *A. ipaensis*) fall into the Bolivia-dominated Clade 4. This clade contains *hypogaea* and *fastigiata* varieties, including the uncommon aequatoriana variety, which is classified (Krapovickas et al. 2007) as *A. hypogaea* subsp. *fastigiata* var. *aequatoriana*. The *aequatoriana* variety is generally not widely used in cultivation outside of the landrace occurrence in these regions in South America. Krapovickas (1995) describes *A. hypogaea* subsp. *hypogaea* var. *hirsute, A. hypogaea* subsp. *fastigiata* var. *peruviana*, and *A. hypogaea* subsp. *fastigiata* var. *aequatoriana* as being important in ancient times, and still important locally, being found in Peruvian markets, for example. These highly reticulated pod types are also seen in multiple archaeological sites on the coast of Peru and Chile, and Argentina (Masur et al. 2018; Krapovickas 1995), as well as in early European herbarium specimens. This pod form is depicted in the royal tombs of Sipán, in northern Peru, dating ca. 250 AD, associated with the Moche culture (Krapovickas 1995; Masur et al. 2018). The peanut form in the necklace, sculpted clearly in gold and silver, is identified by Krapovickas (1995) as *A. hypogaea, subsp. fastigiata var. peruviana.* (Supplementary Figure S15).

The identification of southeastern Bolivia, as the center of origin of cultivated peanut relies on several lines of evidence. Both ancestral diploid species *A. duranensis* and *A. ipaensis* are found close to Villa Montes, in the Province of Tarija (Krapovickas et al. 2009; Krapovickas and Gregory 1994). These species are strongly prostrate, lack flowers on the main stem, have dark green leaves, and small two seeded pods (Krapovickas et al. 2009; Krapovickas and Gregory 1994). Also in Tarija are found a large number of var. *hypogaea* landraces including the archetypal primitive cultivated peanut, “Rastrero colorado de dos granos,” which combines the most primitive characteristics being, a strongly prostrate variety, with dark green leaves, lacking flowers on the main stem, and most importantly, it has two seeded pods with small seeds (Krapovickas et al. 2009; Krapovickas and Gregory 1994). This combination of prostrate habit and small seeds is very rare. The sample studied here did not include Rastrero colorado de dos granos, but it is notable that Clade 4 includes other landraces with these Tarija primitive characteristics. These include Sara Maní (PI 468280), from nearby Cochabamba Province, which has pods that are very similar to Rastrero colorado de dos granos, except with a slightly less prominent beak (Krapovickas et al. 2009). Also very notably, Clade 4 contains all nine of the smallest seeded var. *hypogaea* types (prostrate and lacking flowers on the main stem): PI 336978, PI 442768, PI 210831, PI 497342, PI 331337, PI 471986, PI 288210, PI 221068, and PI 468280.

### Network and Structure analysis

To further define subpopulations and the genetic relatedness among accessions, we performed a structure and network analysis (Figure 3). At K = 5, accessions were assigned into groups that corresponded with phylogenetic and network assignments in Figures 1 and 2.

Clusters 1 and 2 had the most membership and cluster 3 the least (166, 164, 19) (Supplementary Files S10 and S11). Based on the global ancestry estimates on all genomic SNP sequences (Raj et al. 2014), accessions were colored in accordance with cluster assignment. An accession that could not be assigned to a definitive cluster was painted admixed with colors representative of each cluster with which it proportionally shared genomic sequences.

Overall, 240 accessions were exclusively assigned to a single group and 278 were assigned, in admixed proportions, to two or three groups: with 221 assigned to two, and 57 to three clusters. Of the 12 check cultivars genotyped, eight were assigned to cluster 4, along with Tifrunner. Of these, only Jupiter was exclusively assigned to a single cluster with the remaining seven, including Tifrunner, sharing admixed proportions with more than one cluster. Fuhuasheng and Shitouqi were assigned to cluster 2, same as cultivars Olin and Tamnut OL 06. The synthetic tetraploid sequence “duranensis_ipaensis” was assigned to cluster 5 - the only cluster without any cultivar assigned.

Clustering accessions via a phylogenetic analysis is overly simplistic as it suggests a one-dimensional source for sequence similarity or dissimilarity between a pair of accessions. Network analysis provides a more representative and explanatory relationship between given accessions. In Figure 2, accessions with similar sequence characteristics cluster near each other in the network. The further apart accessions are in the network, the more different they are in sequence characteristics (Figure 2). Four main clusters were defined representing accessions that were more similar to each other and distinct from those in other clusters. Even though most accessions cluster in close correspondence to phylogenetic cluster definitions, exceptions show that a bifurcating tree representation of sequence similarity may not represent the true underlying nature of relatedness among accessions.

Overall, we found groups defined on phylogenetic clade membership to correspond with groups defined by structure and network analyses. These groups showed high genetic differentiation. Clade 1 was genetically distinct from Clades 2, 3, 3.2 and 4 (F_STs_ : 0.74, 0.75, 0.67, 0.51). Clades 3 and 3.2 were not much different from each other (F_ST_ 0.22). Clade 3.2 was also not strongly distinct from Clade 2 (F_ST_ 0.3). Genetic clustering via PCA confirmed the main groups as distinct clusters (Figure 4A, Figure 5A).

#### Genetic diversity correlates with subspecies and botanical types

Principal Coordinates 1 and 2 (PC1 and PC2), which together explained 59.75 % of the genetic variation in the collection, differentiated between the two subspecies and corresponding botanical varieties. PC1 separated ssp. *hypogaea* from ssp. *fastigiata*, while PC2 delineated between the two ssp. *fastigiata* varieties; separating var. *fastigiata* from var. *vulgaris*. PC1 also corresponded with Virginia and Runner type accessions while PC2 separated Spanish types from Valencia types (Figures 4B,C).

These results suggest a pattern, consistent with the biology of subspecies and botanical variety classification, as the most important correlates of the genetic diversity in the collection. Previous studies using a subset of this collection, the mini core, have suggested the presence of between four to five sub-populations (Otyama et al. 2019; Wang et al. 2011; Belamkar et al. 2011). These results recapitulate and add support to these findings, further linking biology to the landscape of genetic stratification in the U.S. peanut core collection.

Growth form and pod shape did not correspond well with PCA even though both traits are key determinants of agronomic type classification, and, by extension, subspecies groups (Figures 4D,E). Pod shape considers the constriction, reticulation, and the number of seeds per pod to define five main groups: *vulgaris, fastigiata, peruviana, hypogaea* and *hirsuta*. Spanish and Valencia types are classified as “bunch” for their upright growth form while Virginia and Runner types are classified as “runners” for their prostrate (flat) growth form. Several Virginia varieties are also classified as “decumbent”, for their intermediate growth form between “runner” and “bunch” (Pittman 1995). The lack of a clear correspondence between growth form and pod shape with genetic diversity, begs for more studies with special emphasis on accurate phenotyping, to help establish their contribution to genetic stratification and diversity in peanut collections.

#### Genetic differentiation among groups (F_ST_ fixation index)

The genetic difference between varieties belonging to contrasting subspecies was relatively high. Accessions classified as var. *vulgaris* appeared genetically distinct from those classified as var. hypogaea with F_ST_ 0.59. The difference was comparatively low for varieties of the same subspecies, var. *vulgaris* and var. *fastigiata* accessions, F_ST_ 0.198 (Figure 5B). This provides clear evidence for the genetic distinction between subspecies and corresponding botanical varieties.

Interestingly, a comparison between var. *aequatoriana* and var. *fastigiata* showed a surprisingly high level of differentiation, F_ST_ 0.40. Since both varieties are classified as ssp. *fastigiata*, genetic differentiation was expected to be much smaller. Contrastingly, we observed low genetic separation for an inter-subspecies comparison between var. *aequatoriana* and var. *hypogaea*, F_ST_ 0.2 (Figure 5B). This result suggests a possible misclassification of var. *aequatoriana* accessions, which share greater similarity to ssp. *hypogaea* than the ssp. *fastigiata* group to which they are assigned. Evidence for misclassification was first suggested by (He and Prakash 2001; Raina et al. 2001; Ferguson et al. 2004; Tallury et al. 2005; Freitas et al. 2007; Cuc et al. 2008) and later alluded to by (Bertioli et al. 2011). However, like their studies, this present analysis suffers from a low number of var. *aequatoriana* accessions. Additionally, only 159 samples representing accessions in the core, have been classified. Of these, 114 are classified as var. *fastigiata*, 43 as var. *hypogaea* and two as var. *aequatoriana* (Data source: GRIN). Since within-population diversity has been shown to affect F_ST_ as an estimate of genetic differentiation among populations (Hedrick 1999; Bird et al. 2011), we recommend cautious interpretation of these results, especially where they conflict with known peanut biology.

Market types, Spanish and Virginia, showed evidence of genetic differentiation (F_ST_ 0.4), as did Valencia and Virginia (F_ST_ 0.4), and Valencia and Spanish (F_ST_ 0.3) (Figure 5C). Indeed, accessions marked as “mixed” showed low pairwise genetic differentiation with main groups – as would be expected from a phenotypically ambiguous group. As expected, Runner accessions were more similar to Virginia accessions (F_ST_ 0.027) compared to Valencia (F_ST_ 0.29), and Spanish types (F_ST_ 0.26) (Figure 5C). Classification studies place Valencia and Spanish types under the same subspecies, ssp. *fastigiata*, but different botanical varieties - var. *fastigiata* and var. *vulgaris*, respectively. Virginia types are classified under a different subspecies altogether - ssp. *hypogaea* var. *hypogaea*. This result supports Runner types as a hybrid between the two peanut subspecies as classified by Krapovickas (1969).

Non-distinct phenotypes like pairwise comparisons of growth forms: “spreading-bunch”, “spreading”, “bunch” and “mixed”, which are affected by environmental conditions, resulted in less pronounced genetic separation among groups. The contrast was true with phenotypically distinct groups for pairwise comparisons between growth forms: “spreading” and “prostrate” (F_ST_ 0.55), “spreading” and “erect” (F_ST_ 0.39), “spreading-bunch” and “erect” (F_ST_ 0.28) (Figure 5D). This suggests a good prediction of phenotypic diversity by genetic variation. Groups defined under pod shape were not distinct from each other suggesting phenotypic ambiguity in these classes (Figure 5E).

Collectively, these results suggest a level of stratification that is consistent with subspecies groups and botanical variety classification. Overall, we found accessions were similar within botanical varieties and subspecies groups, but genetic separation increased evidently between group comparisons. This carries important implications for studies using this collection for genetic associations. Treating the collection as a homogenous group may obscure association results and if not properly accounted for, population stratification may cause studies to fail due to lack of significant results or overwhelming false-positive signals.

**Geographic origin does** not **generally** correspond with genetic diversity

On the whole, the country of seed origin was not an important contributor to structure in the collection. There was little genetic differentiation between peanuts based on where seed was originally collected. African and North American accessions appeared genetically similar (F_ST_ 0.02), as did Asian and African accessions (F_ST_ 0.01) (Figure 5F).

We also found the country of seed origin to be a poor correlate of genetic structure, even though the core collection is predominated by accessions from South America and Africa, which together make up 74.6% of the entire collection. The peanuts collected from Bolivia and South America were not so distinct as to cluster around a recognizable pattern or separate from those collected from other continents. This may suggest that not many independent mutations have arisen in the different continental subgroups to cause significant genetic separation. It is also known that peanuts had completely differentiated into subspecies and botanical varieties prior to being dispersed from their center of origin by early explorers and traders (Simpson et al. 2001).

### The mini core is representative of the genetic diversity in the core collection

The mini core collection was created to further define a small manageable sub collection representative of the diversity in the germplasm collection. The need was driven by a reliance on low-throughput markers, like RFLPs and SSRs, which are difficult and costly to assay in large collections and some agronomic traits being quite difficult and costly to measure (Holbrook and Dong 2005). We used genetic clustering via PCA to define how well the mini core represents the diversity in the core collection.

Results show remarkable representation spanning the entire spread of the genetic diversity in the core collection (Figure 4F). Thus, clustering on select morphological characteristics followed by sampling within defined clusters likely resulted in the selection of a well representative set. The main weakness of the mini core is its relatively small size (94 available accessions), which weakens the ability to identify novel marker-trait associations in genome-wide association studies (Otyama et al. 2019). However, the mini core collection has proven to be of much utility for identifying germplasm with desirable characteristics for breeding pipelines and for verifying identified marker-trait associations (Holbrook and Dong 2005; Dean et al. 2009; Wang et al. 2011).

### Subgenome exchanges are a significant source of diversity in tetraploid peanut

An enduring puzzle regarding peanut evolution is that the diversity in the crop appears to have arisen quickly, from a severe genetic bottleneck at the time of the tetraploidization event roughly 10,000 years ago, likely involving a rare, single plant in an early horticulturalist’s garden (Bertioli et al. 2019). The diploid progenitors, *A. duranensis* and *A. ipaensis*, separated approximately 2 million years ago (Bertioli et al. 2016), and the best evidence is that the mergers of these diploids has occurred only once in pre-modern times (Bertioli et al. 2016; Bertioli et al. 2019). To put the question simply: how did so much genotypic and phenotypic diversity arise in modern peanut varieties?

One source of the diversity was identified by (Bertioli et al. 2019), with the reporting of the high-quality Tifrunner genome sequence. Specifically, exchanges between corresponding chromosomes of the A and B genomes were seen - on scales both small (on the gene-scale), and large (on the scale of multiple megabases, at chromosome ends). We used the genotyping data from the current project to independently assess the patterns of subgenome exchanges.

In the variation data from the Affymetrix array, we found evidence of both widespread small-scale exchanges between subgenomes, and apparent large-scale “invasions” of one subgenome to the other. These patterns are evident in Figure 7, shown in red, whereas gray indicates loci where subgenome exchange either was not observed or there was insufficient evidence regarding exchange. One pattern to note is that different accessions show different patterns. Each of the 16 diverse accessions used for comparison is represented along a vertical slice next to each chromosome. At high resolution, many between-accession differences can be seen - for example, at the top of A01, where the first two accessions show an exchange, and the middle accessions do not. Also noteworthy are regions that were reported, in the Tifrunner genome paper, to show invasion (and replacement) of one subgenome by the other. In these locations (marked in green along the chromosome backbones), most alleles are either all red, indicating that the chromosomal segment was contributed by the other subgenome; or all gray, indicating that the chromosomal segment was contributed by the “cis” subgenome. This is evident at the top of A05 and B05, for example.

Of the 10,829 SNP positions for which it was possible to evaluate subgenome exchanges (as data was present for all tested lines), there was evidence of exchanges in at least one accession for 1,068 positions (9.8%). This is likely a highly conservative estimate, as many positions are ambiguous with respect to subgenome origin - for example, when the reference SNPs from the diploids may be from the other allele (not represented in the genome sequence).

Our interpretation is that a substantial fraction (>10%) of alleles have arisen through subgenome exchanges; and further, that these exchanges appear to be ongoing, as there are numerous differences between accessions, in the subgenome allele status at a given locus.

## Conclusions

Genotype data for each accession in the U.S. peanut core collection will benefit peanut breeders in multiple ways: providing SNP data for use in marker-trait association studies to identify SNPs associated with important traits, describing the population structure of the core, and enabling breeders to work with smaller groups of accessions by selection through both phenotypic and genotypic characteristics. A probable ancestral genotypic group is identified, with most such accessions still coming from near the geographical origin of tetraploid peanut. The data also provides information about the ongoing rapid changes in the peanut genome through subgenome exchanges, and supports theories about the origin, early cultivation, and dispersion of peanut throughout the world.

## Acknowledgements

This project was supported by the Agriculture and Food Research Initiative Competitive Grant no. 2018-67013-28138 co-funded by the USDA National Institute of Food and Agriculture and the National Peanut Board. Mention of trade names or commercial products in this publication is solely for the purpose of providing specific information and does not imply recommendation or endorsement by the U.S. Department of Agriculture. USDA is an equal opportunity provider and Employer. We thank Dr. Josh Clevenger for insights regarding subgenome exchanges, Dr. Guillermo Seijo for information regarding the origins of cultivated peanut, Dr. H. Eric R. Olsen regarding Portuguese and Spanish colonial trade of peanut, Dr. Shyam Tallury for assistance with germplasm, and Dr. Naveen Puppala for permitting the use of NuMex-01 before its public release.

## Literature Cited

Altschul, S.F., W. Gish, W. Miller, E.W. Myers, and D.J. Lipman, 1990 Basic local alignment search tool. Journal of molecular biology 215 (3):403–410.

Anon., 2000 Release of ‘Jupiter’ peanut. Oklahoma State University, Oklahoma Agricultural Experimental Station, USA.

Baring, M.R., Y. Lopez, C.E. Simpson, J.M. Cason, J. Ayers et al., 2006 Registration of ‘Tamnut OL06’ peanut. Crop Science 46 (6):2720–2721.

Baring, M.R., C.E. Simpson, M.D. Burow, J.M. Cason, and J. Ayers, 2013 Registration of ‘Tamrun OL11’ Peanut. Journal of Plant Registrations 7 (2):154.

Belamkar, V., M.G. Selvaraj, J.L. Ayers, P.R. Payton, N. Puppala et al., 2011 A first insight into population structure and linkage disequilibrium in the US peanut minicore collection. Genetica 139 (4):411.

Bertioli, D.J., S.B. Cannon, L. Froenicke, G. Huang, A.D. Farmer et al., 2016 The genome sequences of Arachis duranensis and Arachis ipaensis, the diploid ancestors of cultivated peanut. Nature Genetics 48:438.

Bertioli, D.J., J. Jenkins, J. Clevenger, O. Dudchenko, D. Gao et al., 2019 The genome sequence of segmental allotetraploid peanut Arachis hypogaea. Nature genetics 51 (5):877–884.

Bertioli, D.J., G. Seijo, F.O. Freitas, J.F. Valls, S.C. Leal-Bertioli et al., 2011 An overview of peanut and its wild relatives. Plant Genetic Resources 9 (1):134–149.

Bird, C.E., S.A. Karl, P.E. Smouse, and R.J. Toonen, 2011 Detecting and measuring genetic differentiation. Phylogeography and population genetics in Crustacea 19 (3):1-55.

Branch, W., 2007a Registration of ‘Georgia-06G’peanut. Journal of Plant Registrations 1 (2):120–120.

Branch, W.D., 2007b Registration of ‘Georgia-06G’ Peanut. Journal of Plant Registrations 1 (2):120–120.

Chamberlin, K., R. Bennett, J. Damicone, C. Godsey, H. Melouk et al., 2015 Registration of ‘OLe’peanut. Journal of Plant Registrations 9 (2):154–158.

Chen, X., Q. Lu, H. Liu, J. Zhang, Y. Hong et al., 2019 Sequencing of cultivated peanut, Arachis hypogaea, yields insights into genome evolution and oil improvement. Molecular plant 12 (7):920–934.

Clevenger, J.P., W. Korani, P. Ozias-Akins, and S. Jackson, 2018 Haplotype-based genotyping in polyploids. Frontiers in plant science 9:564.

Cuc, L.M., E.S. Mace, J.H. Crouch, V.D. Quang, T.D. Long et al., 2008 Isolation and characterization of novel microsatellite markers and their application for diversity assessment in cultivated groundnut (Arachis hypogaea). BMC plant biology 8 (1):55.

Dash, S., C.E.K. S., K.S. R, F.A. D., and C.S. B., 2016 PeanutBase and Other Bioinformatic Resources for Peanut.

Dean, L., K. Hendrix, C. Holbrook, and T. Sanders, 2009 Content of some nutrients in the core of the core of the peanut germplasm collection. Peanut Science 36 (2):104–120.

Ferguson, M., P. Bramel, and S. Chandra, 2004 Gene diversity among botanical varieties in peanut (Arachis hypogaea L.). Crop Science 44 (5):1847–1854.

Francis, R.M., 2017 pophelper: an R package and web app to analyse and visualize population structure. Molecular Ecology Resources 17 (1):27–32.

Freitas, F., M. Moretzsohn, and J. Valls, 2007 Genetic variability of Brazilian Indian landraces of Arachis hypogaea L. Embrapa Recursos Genéticos e Biotecnologia-Artigo em periódico indexado (ALICE).

Gorbet, D., and B. Tillman, 2009 Registration of ‘Florida-07’peanut. Journal of Plant Registrations 3 (1):14–18.

Goudet, J., 2005 Hierfstat, a package for R to compute and test hierarchical FIstatistics. Molecular Ecology Notes 5 (1):184–186.

He, G., and C. Prakash, 2001 Evaluation of genetic relationships among botanical varieties of cultivated peanut (Arachis hypogaea L.) using AFLP markers. Genetic Resources and Crop Evolution 48 (4):347–352.

Hedrick, P.W., 1999 Perspective: highly variable loci and their interpretation in evolution and conservation. evolution 53 (2):313–318.

Holbrook, C.C., W.F. Anderson, and R.N. Pittman, 1993 Selection of a Core Collection from the U.S. Germplasm Collection of Peanut. Crop Science 33 (4):859–861.

Holbrook, C.C., and W. Dong, 2005 Development and Evaluation of a Mini Core Collection for the U.S. Peanut Germplasm Collection. Crop Science 45 (4):1540.

Holbrook, C.C., P. Timper, A.K. Culbreath, and C.K. Kvien, 2008 Registration of ‘Tifguard’peanut. Journal of Plant Registrations 2 (2):92–94.

Huson, D.H., and D. Bryant, 2006 Application of Phylogenetic Networks in Evolutionary Studies. Molecular Biology and Evolution 23 (2):254–267.

Isleib, T., S. Milla-Lewis, H. Pattee, S. C Copeland, C. Zuleta et al., 2011 Registration of ‘Bailey’ Peanut.

Kassambara, A., 2016 ggcorrplot: Visualization of a Correlation Matrix using’ggplot2’. R package version 0.1 1.

Korani, W., J.P. Clevenger, Y. Chu, and P. Ozias-Akins, 2019 Machine Learning as an Effective Method for Identifying True Single Nucleotide Polymorphisms in Polyploid Plants. Plant Genome 12 (1).

Krapovickas, A., 1969 The origin. variability and spread of the groundnut (Arachis hypogaea).

Krapovickas, A., 1995 El origen y dispersión de las variedades del maní.

Krapovickas, A., and W.C. Gregory, 1994 Taxonomia del genero Arachis (Leguminosae). Bonplandia VIII:1–187.

Krapovickas, A., W.C. Gregory, D.E. Williams, and C.E. Simpson, 2007 Taxonomy of the genus Arachis (Leguminosae). Bonplandia 16:7–205.

Krapovickas, A., and R.O. Vanni, 2009 El maní de Llullaillaco. Bonplandia 18 (1):51–55.

Krapovickas, A., R.O. Vanni, J.R. Pietrarelli, D.E. Williams, and C.E. Simpson, 2009 Las razas de maní de Bolivia. Bonplandia 1:95–189.

Masur, L.J., J.-F. Millaire, and M. Blake, 2018 Peanuts and Power in the Andes: The Social Archaeology of Plant Remains from the Virú Valley, Peru. Journal of Ethnobiology 38 (4):589–609.

Melouk, H.A., K. Chamberlin, C.B. Godsey, J. Damicone, M.D. Burow et al., 2013 Registration of ‘Red River Runner’peanut. Journal of Plant Registrations 7 (1):22–25.

Otyama, P.I., A. Wilkey, R. Kulkarni, T. Assefa, Y. Chu et al., 2019 Evaluation of linkage disequilibrium, population structure, and genetic diversity in the US peanut mini core collection. BMC genomics 20 (1):481.

Pittman, R.N., 1995 United States peanut descriptors. ARS (USA).

Price, M.N., P.S. Dehal, and A.P. Arkin, 2010 FastTree 2–approximately maximum-likelihood trees for large alignments. PloS one 5 (3).

Puppala, N., and S.P. Tallury, 2014 Registration of ‘NuMex 01’ High Oleic Valencia Peanut. Journal of Plant Registrations 8 (2):127.

Raina, S., V. Rani, T. Kojima, Y. Ogihara, K. Singh et al., 2001 RAPD and ISSR fingerprints as useful genetic markers for analysis of genetic diversity, varietal identification, and phylogenetic relationships in peanut (Arachis hypogaea) cultivars and wild species. Genome 44 (5):763–772.

Raj, A., M. Stephens, and J.K. Pritchard, 2014 fastSTRUCTURE: Variational Inference of Population Structure in Large SNP Data Sets. Genetics 197 (2):573–589.

Rognes, T., T. Flouri, B. Nichols, C. Quince, and F. Mahé, 2016 VSEARCH: a versatile open source tool for metagenomics. PeerJ 4:e2584.

Simpson, C., M. Baring, A. Schubert, H. Melouk, Y. Lopez et al., 2003 Registration of’OLin’peanut. Crop Science 43 (5):1880–1882.

Simpson, C., A. Krapovickas, and J. Valls, 2001 History of Arachis including evidence of A. hypogaea L. progenitors. Peanut Science 28 (2):78–80.

Tallury, S., K. Hilu, S. Milla, S. Friend, M. Alsaghir et al., 2005 Genomic affinities in Arachis section Arachis (Fabaceae): molecular and cytogenetic evidence. Theoretical and Applied Genetics 111 (7):1229–1237.

Tillman, B., and D. Gorbet, 2015 Registration of ‘FloRun ‘107’’peanut. Journal of Plant Registrations 9 (2):162–167.

Wang, M.L., S. Sukumaran, N.A. Barkley, Z. Chen, C.Y. Chen et al., 2011 Population structure and marker–trait association analysis of the US peanut (Arachis hypogaea L.) mini-core collection. Theoretical and Applied Genetics 123 (8):1307–1317.

Weir, B.S., and C.C. Cockerham, 1984 Estimating FIstatistics for the analysis of population structure. evolution 38 (6):1358–1370.

Zheng, X., D. Levine, J. Shen, S.M. Gogarten, C. Laurie et al., 2012 A high-performance computing toolset for relatedness and principal component analysis of SNP data. Bioinformatics 28 (24):3326–3328.

Zhuang, W., H. Chen, M. Yang, J. Wang, M.K. Pandey et al., 2019 The genome of cultivated peanut provides insight into legume karyotypes, polyploid evolution and crop domestication. Nature Genetics 51 (5):865–876.

